# Rab2 is a potent new target for enhancing autophagy in the treatment of Parkinson’s disease

**DOI:** 10.1101/2020.08.30.274050

**Authors:** Janka Szinyákovics, Eszter Kiss, Fanni Keresztes, Tibor Vellai, Tibor Kovács

## Abstract

Macroautophagy is a lysosomal-dependent degradational pathway of eukaryotic cells, during which toxic, unnecessary, and damaged intracellular components are broken down. Autophagic activity declines with age, and this change could contribute to the accumulation of intracellular damage at advanced ages, causing cells to lose their functionality and vitality. This could be particularly problematic in post-mitotic cells include neurons, the mass destruction of which leads to different neurodegenerative diseases.

We aim to discover new regulation points where autophagy could be specifically activated, and test these potential drug targets in Drosophila neurodegenerative disease models. One possible way to activate autophagy is through the enhancement of autophagosome-lysosome fusion to become autolysosome. This fusion is regulated by HOPS (homotypic fusion and protein sorting) and SNARE (Snap receptor) complexes. The HOPS complex forms a bridge between lysosome and autophagosome with the assistance of small GTPase Rab (Ras-associated binding) proteins. Thus, Rab proteins are essential for autolysosome maturation, and among Rab proteins, Rab2 is required for the degradation of autophagic cargo.

Our results revealed that GTP-locked (constitutively active) Rab2 (Rab2 CA) expression reduces the levels of the autophagic substrate p62/Ref2P in dopaminergic neurons, and improved the climbing ability of animals during aging. The expression of Rab2 CA also increased lifespan in a Parkinson’s disease model (human mutant alpha-synuclein [A53T] overexpressed animals). In these animals, Rab2 CA expression significantly increased autophagic degradation as compared to control. These results may reveal a new, more specific drug target for autophagic activation treating today’s incurable neurodegenerative diseases.

## Introduction

In eukaryotic cells, there are two main forms of protein degradation. One of them is the ubiquitin-proteosome system, which degrades proteins that are labelled by a multiubiquitin signal^1^.The other is autophagy, which can breakdown both damaged proteins and protein aggregates. In addition to protein degradation, autophagy can eliminate whole organelles^2^. Autophagy is a lysosomal degradation pathway, which has three main types depending on how the cargo is delivered into the lysosomal compartment: microautophagy, chaperone-mediated autophagy (CMA) and macroautophagy.

During macroautophagy (hereafter referred to as autophagy), a double-membrane structure called phagophore or isolation membrane forms in the cytoplasm around damaged or superfluous proteins and organelles, which if not degraded are toxic to the cell. This process is called vesicle nucleation. With the closure of phagophore membrane, the structure becomes an autophagosome, which can fuse with a lysosome to create an autolysosome or with an endosome to make an amphisome, which then also fuses with a lysosome. During lysosomal degradation, the enclosed cargo is degraded by acidic lysosomal hydrolases^3^. Because damaged proteins and organelles interfere normal cellular functions, autophagy has a crucial cytoprotective role in eliminating such components. In addition, it has also important roles in macromolecule recycling and in providing energy for cellular processes^4^.

Autophagy is essential for maintaining cellular homeostasis. However, as it was demonstrated in several genetic models, the capacity of autophagy declines with age, leading to a significant accumulation of cellular damage at advanced ages^5-6^. Thus, the process plays an important role in slowing down the rate at which cells age^7-8-9^. For example, in the nematode *Caenorhabditis elegans*, loss-of-function mutations in autophagy genes (*atg*), including *atg1/unc-51* (uncoordinated), *atg18* and *atg6/bec-1* (beclin 1 homolog), cause progeria and lifespan shortening^6-10^. Furthermore, enhancing autophagy by overexpressing *atg8* or *atg5* genes extends lifespan in *Drosophila melanogaster and Mus musculus*^11-12^. A decrease in the capacity of autophagy is therefore a significant problem for post-mitotic cells like neurons, which lose the ability to divide, so it is not possible for them to dilute damaged constituents by cell division^13^. Decline in autophagic degradation often leads to the incidence of various neurodegenerative disorders^14-15^.

In neurons, autophagosomes are formed at the distal end of axons (right at the presynaptic membrane) and transported along microtubules by motor proteins, such as dyneins, to the some (ret-rograde transport)^16-17^. To recruit dynein, it is essential for autophagosomes to fuse with late endosomes, resulting in structures called amphisomes^18^. During transport to the soma, co-localization of key autophagy proteins LC3 (Light chain 3B) and LAMP1 (Lysosomal-associated membrane protein 1) occurs, and the acidification of autophagic structures are also evident, suggesting the fusion of amphisomes with lysosomes^19^. Taken together, these observations suggest that in neurons autophagosomes undergo a spatially defined maturation process, which requires endocytosis and retrograde transport^19^.

Autophagy dysfunction is involved in the pathology in several neurodegenerative conditions^20^. In Parkinson’s disease (PD), for example, mutant α-synuclein, which is normally broken down by CMA, binds strongly to LAMP2A transporter on the lysosome and is therefore unable to translocate into the lysosome^21^. It also inhibits the translocation of other proteins into the lysosomal lumen, thereby increasing the accumulation of defective cytoplasmic proteins in the cell^21^. Furthermore, mutations in proteins that are involved in PD, including Parkin, PINK1 (PTEN-induced kinase 1) and PARK7 (Parkinson disease protein 7), also affect autophagic degradation. In case of mitochondrial damage, these proteins can ubiquitinite outer mitochondrial membrane proteins. Thus, they degrade defective organelles by selective autophagy called mitophagy. However, in the absence of these proteins, mitophagy is blocked, making the cell more sensitive to oxidative stress. Reactive oxygen species (ROS) released into the cytoplasm eventually lead to neuronal cell loss. Dopaminergic neurons in the midbrain substantia nigra pars compacta are particularly affected by this process^22^.

At present, neurodegenerative diseases represent as largely uncurable human pathologies. This creates a great demand for understanding the molecular basis of such diseases and find new targets for their effective treatment. The risk for developing various neurodegenerative diseases gradually increases with age. Therefore, there is currently an intensive research for small molecules (drug candidates) activating autophagy which can be potentially used to treat neurodegenerative pathologies. Autophagy inducers identified so far largely influence processes that act upstream of autophagy, often leading to undesired side effects. Most of these inducers target TOR (target of rapamycin) kinase, which integrates the cellular information on nutrient and growth factor availability^23^. In yeast, there are two TOR paralogs, TOR1 and TOR2, while in higher eukaryotes there is one TOR protein that participates in two complexes, TORC1 (TOR complex 1) and TORC2 (TOR complex 2)^24^. The function of these TOR paralogs and complexes differs from each other. TOR1 and TORC1 are associated with autophagy, cell growth, protein synthesis, cellular metabolism and cell cycle regulation, while TOR2 and TORC2 regulate the cytoskeletal system and lipid synthesis. Thus, TOR regulates various cellular processes in addition to autophagy, thus manipulating it’s activity can lead to severe effects.

During vesicle nucleation, the VPS34 (Vacuolar protein sorting 34) kinase complex marks vesicles of various origins with PI3P (Phosphatidylinositol 3-phosphate), thereby linking these structures to the autophagic pathway^25^. An antagonist of this process is MTMR14 (myotubularin-related lipid phosphatase) protein, which is capable of dephosphorylating PI3P through it’s phosphatase activity, thereby inhibiting the initiation of autophagy^26^. We previously showed that two autophagy enhancers, AUTEN-67 (autophagy enhancer) and AUTEN-99, exert their autophagy-activating effect through inhibiting MTMR14^26-27-28^. These drug candidates increase autophagy activity in cell cultures and in various model organisms including *Drosophila*, zebrafish and mice^26-28^. In *Drosophila*, both AUTEN-67 and -99 promote longevity and improve the ability of aged animals to climb up on the wall of glass vials. Moreover, these AUTEN molecules significantly lower the amount of toxic protein aggregates found in *Drosophila* models of PD and Huntington’s disease (HD), thereby increasing the survival of affected neurons^27-28^.

Both AUTEN-67 and 99 activate early stages of the autophagic process, phagophore and autophagosome formation, at which the type III PtdIns3 kinase complex functions. However, some studies suggest that inadequate degradation of the autophagosomal content may contribute to the formation and accumulation of toxic proteins causing neurodegrenerative processes^29^. This prompted us to identify novel target proteins, the modulation of which can promote acidic degradation after autophagosome formation. One possible way is to enhance the autophagosome-lysosome fusion to promote autolysosome formation. The fusion process is regulated by HOPS (homotypic fusion and protein sorting) and SNARE (Snap receptor) complexes, which use small GTPase proteins such as Rab (Ras-associated binding) to link together the two vesicle structures. Rab proteins have key functions in membrane trafficking, and involved in membrane docking, recruitment of other proteins and membrane fusion^30^. As a molecular switch, the GTP/GDP-binding of Rab strongly influences the binding of effectors to the autophagic membrane. Whether Rab binds to GTP or GDP is dependent on GEFs (guanine nucleotide exchange factors) enzymes that promote GTP-binding and conversion of Rab into an activated, membrane-bound state^31^. Rab is inactivated by GAPs (GTPase-activating proteins) enzymes, which support the hydrolysis of GTP to GDP, resulting in an inactivated, cytosolic form of Rab. Rab2 and Rab7 proteins play an important role in autophagosome maturation^32-33^. Studies on yeast and *Drosophila* found that defective Rab7 or Rab2 cause the accumulation of autophagosomes^34-33^.

In our study we examined the effect of overexpressing constitutively (GTP binded) active Rab2 (named Rab2 CA) in the nervous system of *Drosophila melanogaster*. Our aim was to test whether Rab2 CA overexpression in *Drosophila* has a beneficial effect. We also inspected Rab2 CA overexpression in Parkinson diseased backround in hope for finding a cure for the sideeffects of the disease.

## Materials and Methods

### Animals and Lifespan Assays

We used the UAS-Gal4 system to express UAS-Rab2-CA (constitutively active; BDSC:9760) specifically in dopaminergic neurons (using PleGal4;BDSC:8848; see Fig. 1A) or in dopaminergic and serotonergic neurons (with DdcGal4; BDSC:7010; see Fig. 2B’). We also used UAS-Rab2-WT (BDSC:23246) flies to compare the overexpression of wild-type Rab2 to the overexpression of a constitutively active form of Rab2 (Rab2 CA), and UAS-Luc to compensate 2 Gal4 sequences in a PD model. Animals were kept at 29°C, and only females were studied to avoid differences in sex. During lifespan assays, we measured the number of dead flies every day, and transferred the living animals into new tubes every second day.

**1. Figure.**
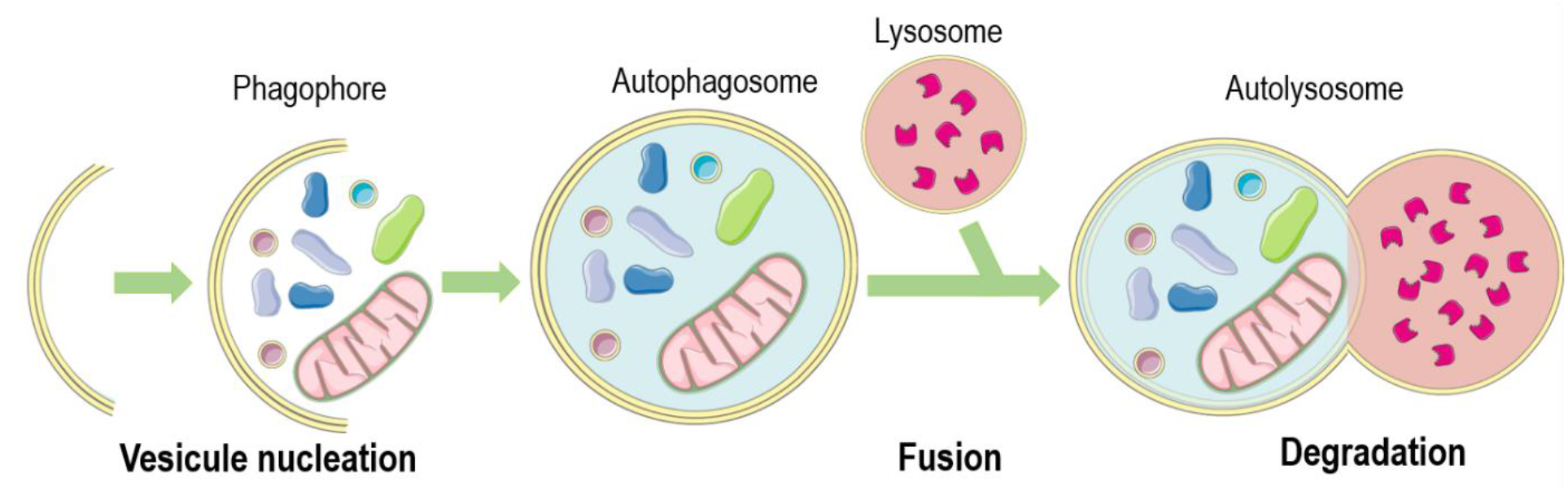
The mechanism of macroautophagy: Macroautophagy starts by forming a double membrane structure called phagophore in the cytosol around proteins and organelles that are destined to be degraded. When membrane growing becomes completed, a vacuole called autophagosome is formed which then fuses with a lysosome to generate an autolysosome, in which the enclosed structures are degraded by acidic hydrolases.

**Figure 2.**
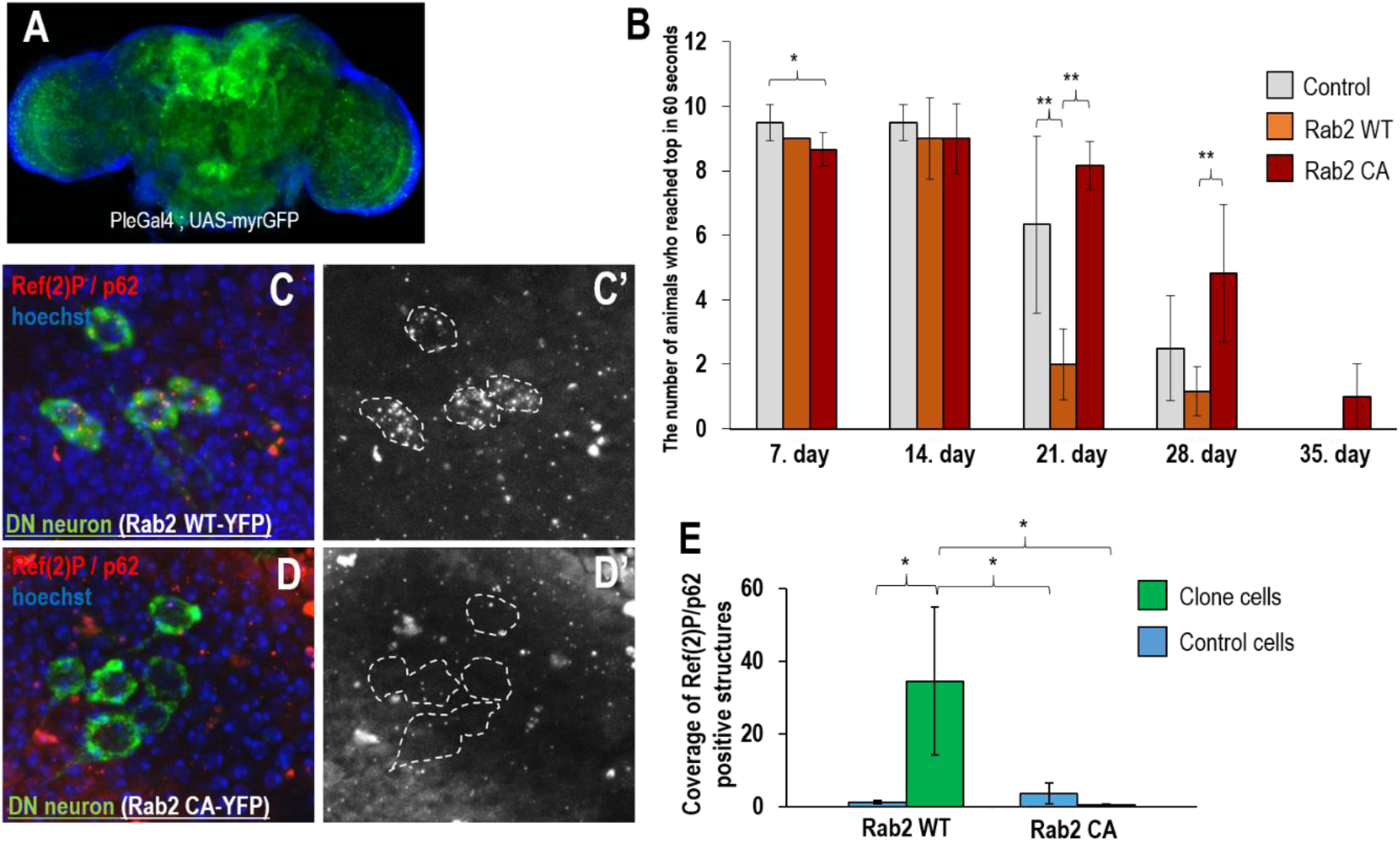
Constitutively active Rab2 in dopaminerg neurons improves climbing ability in elderly flies. (**A**)The regions of Drosophila nervous system where pleGal4 is expressed, showed by an UAS-myrGFP fluorenscent reporter (green). (**B**) Climbing ability of flies at different adult stages. 25 cm long glass vials were used to monitor the ability of animals to climb on the wall of test tubes. Overexpression of Rab2 CA in dopaminergic neurons (DNs) improves the climbing ability of aged flies, while overexpression of Rab2 WT is not effective in this test. Animals were tested for 60 sec after shaking them down. (**C-E**) DNs expressing Rab2 WT (**C-C’**) appear to contain more Ref2P/p62-positive structures as compared to control cells (**E**). Raf2P/p62 is an autophagic sub-strate, hence its increased levels indicate defects in autophagic degradation. Overexpression of Rab2 CA is associated with fewer amounts of Ref2P/p62 positive structures (**D-D’**). This indicates activated autophagic degradation in the affected DNs (**E)**. Ref(2)P proteins were visualized by anti-Ref2P/p62 antibody staining. Bars represent ±S.D., *: p<0.05, **: p<0.01; statistical analysis. In panels **A, C** and **D**, blue signal indicates nuclei (Hoechst staining).

### Climbing Assay

After sorting animals into climbing tubes with anesthesia, we provided them an hour to regenerate from anesthesia before the test. In this test, we used the negative geotaxis reflex of animals. We used 25 cm long glass vials to monitor the ability of animals to climb on the wall of these test tubes. We gave the animals 60 seconds to climb and we repeted the measurement 3 times.

### Microscopy

Fluorescent images were captured with a Zeiss Axioimager Z1 upright microscope (with objectives Plan-NeoFluar 10x 0.3 NA, Plan-NeoFluar 40x 0.75 NA and Plan-Apochromat 63×1.4 NA) equipped with ApoTome2. We used AxioVision 4.82 and Jmage J 1.52c softwares to examine and evaluate data.

### Western Blotting

Protein samples of flies stem from female adult heads. 15 μ l samples were loaded on a 4-20% SDS-PAGE and blotted onto Immobilon P PVDF membrane (Millipore, IPVH00010). Next blocking with 0.5% blocking reagent (Roche, 1096176) in PBS containing 0.1% Tween 20. Membranes were probed with the following antibodies: anti-Ref(2)P/p62 (rabbit, 1:2500)^35^, alpha-Tub84B (mouse, 1:2500, Sigma, T6199), anti-Atg8a (rabbit, 1:2500)^36^. The secondary antibodies anti-rabbit IgG alkaline phosphatase (1:1000, Sigma, A3687), anti-mouse IgG alkaline phosphatase (1:1000, Sigma, A8438) were visualized by NBT-BCIP solution (Sigma, 72091).

### Statistical Analysis

For statistical analysis of climbing assays, lifespan measurements (mean lifespan) and fluorescence microscopy, results were determined by using R Studio (Version 3.4.3). Distribution of samples (normal or not) was tested with Lilliefors-test. If it was normal, F-test was performed to compare variances. In cases when variances were equal, two-samples t-test was used, otherwise t-test for unequal variances was applied. In case of non-normal distribution, Mann-Whitney U-test was performed. For life span curve statistics, the logrank (Mantel-Cox) method was used, calculated with the SPSS17.0 program.

## Results

### The activation of Rab2 has a positive effect in older flies

Dopaminergic neurons (DNs) produces the neurotransmitter dopamine, the loss of which can lead to the development of PD. Here we examined the effect of overexpressing wild-type Rab2 (Rab2 WT) and constitutively (GTP-bounded) active Rab2 (Rab2 CA) in DNs in the nervous system of *Drosophila*. We found that in aged animals Rab2 CA improves the ability of animals to climb as compared to both Rab2 WT and control (Fig. 2B). Therefore, activated Rab2 (Rab2 CA) exerts a beneficial effect in old flies. In contrast, overexpression of Rab2 WT appeared to be more harmful than favorable on the climbing ability of animals (Fig. 2B). This effect may result from the fact that the accumulation of the bound form of GDP inhibits autophagy. This is supported by our finding that Ref(2)P/p62, which serves as a substrate for autophagic degradation^35^, accumulated in DNs of samples overexpressing Rab2 WT (Fig. 2C-E).

The amount of Ref(2)P/p62 autophagic substrate protein in the central nervous system of aged animals was examined by immunohistochemistry (Fig. 2C-E). Both Rab2 proteins have a yellow fluorescent protein tag (red foci in green cells on Fig. 2C, D). So, DNs that expressed these Rab2 proteins could be readily detected under fluorescent microscopy. The neighboring cells could be used as a control group in this clonal system. In neurons expressing Rab2 WT, the amount of Ref(2)/p62-positive structures markedly increased as compared to the neighboring cells (Fig. 2C, C’). This indicates decreased levels of autophagy degradation in the affected DNs. In contrast, Rab2 CA-positive neurons contained fewer Ref(2)P/p62-positive structures relative to the neighboring cells, indicating increased levels of autophagic degradation in the affected neurons (Fig. 2D, D’). In sum, this finding suggests that Rab2 CA can enhance the autophagy process in the *Drosophila* nervous system, which has a beneficial effect on neuronal functions in aged flies.

### The activation of Rab2 is beneficial in a model of Parkinson disease

PD can result from an autosomal dominant mutation, such as the A53T point mutation in the human *α-synuclein* gene. The mutant protein is more likely to produce fibrils than the wild-type protein, thereby promoting disease progression. We then used DdcGal4 driver to express Rab2 and A53T in dopaminergic and serotonergic neurons (Fig. 3A). We studied the effect of Rab2 CA overexpression in animals simultaneously overexpressing A53T,. UAS-Luc, UAS-A53T was used to create a fly model of PD. According to our data, Rab2 CA improved climbing ability (Fig. 3C) and increased survival (Fig. 2D) in old flies as compared to the corresponding neurodegrenative control (Luc, A53T). These results also indicate that UAS-Luc;UAS-A53T transgene affects climbing ability and lifspan negatively, successfully generating a PD model.

**Figure 3.**
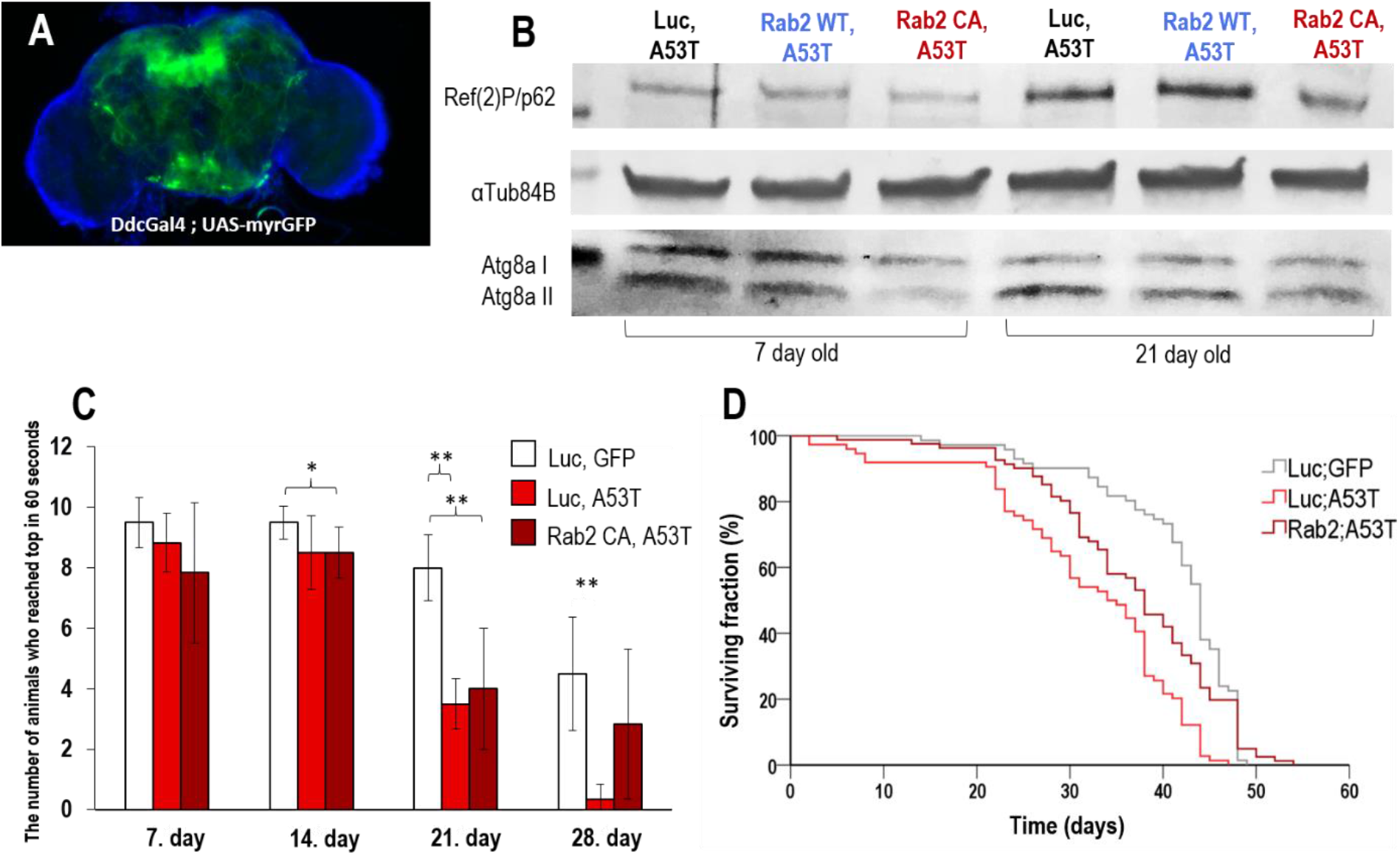
Rab2 CA overexpression interferes with neurodegenerative symptoms in a PD model. (**A**) A DdcGal4 driver and an UAS-myrGFP reporter were co-expressed in dopaminergic and serotonergic neurons (green signals) of the Drosophila brain. Rab2 expression was influenced in these cells. UAS-A53T was used to create flies of PD. (**B**) Proteins were visualized by antibody staining, using a Western-blot assay. Overexpression of Rab2 CA in samples serving as a model of PD lowers levels of Ref2P/p62. This is evident in both age groups. (**C**) Rab2 CA overexpression in a fly model of PD (Luc, A53T) improves climbing ability at late adult stages, as compared to control flies with the same stage. Untreated flies display a decreased ability to climb, indicating that this genetic background serves as a good model for PD. (**D**) Rab2 CA overexpression extends lifespan in a fly model of PD. Untreated animals live shorter than treated ones. Bars represent ±S.D., *: p<0.05, **: p<0.01, statistical analysis.

We finally examined how autophagy activity changes during aging in flies serving as a model for PD. In these animals, Rab2 CA and Rab2 WT were overexpressed. We found that Ref(2)P/p62 levels increase in old animals as compared with young ones (Fig.3/B) in all of them. These results indicate that autophagy becomes compromised in these transgenic animals. Contrary to this, overexpression of Rab2 CA improved autophagic activity as compared to Rab2 WT-overexpressing flies (Fig. 3/B). In Rab2 CA-overexpressing animals, the amount of the lipid-bound Atg8a autophagic membrane protein (Atg8a-II) was reduced, suggesting an efficient degradation of the inner membrane of the autophagosome (Fig.3/B). In summary, activation of Rab2 can activate autophagy in neurons, and improve the survival and cognitive abilities of *Drosophila* serving as a model of PD.

## Discussion

Our results obtained here indicate that overexpression of Rab2 CA reduces the level of the autophagy substrate Ref(2)P/p62protein both in DNs, and improves the movement of aged flies (Fig.2/B-E). In contrast, overexpression of Rab2 WT decreased climbing ability in animals (Fig.2/B). Hence, the bound form of GDP may inhibit autophagy. Overexpression of Rab2 CA in PD animals (flies express the human A53T mutant α-synuclein) extended lifespan and improved climbing ability, as compared to control (Fig.3/C-D). In addition, Rab2 CA increased autophagic degradation relative to control group (Fig.3/B). In the *Drosophila* larval fatbody, Rab2 protein was found to bind to Golgi-derived primary vesicles, while Rab7 was found to bind to autophagosomes or lysosomes^37^. In contrast, in *Drosophila* muscle tissue, Rab2 protein has been described in association with the autophagosome, while Rab7 has been described in association with lysosomes^38,39^. To modulate the activity of Rab2 serving as a potential target for enhancing autophagy, we found that the protein promotes autophagic degradation. So, proteins that inhibit Rab2 serve as potent drug candidate to modulate the autophagic process. These results may be of outstanding significance, as our goal is to find a new, specific autophagy activating targets, providing new drug candidates for the effective treatment of neurodegenerative diseases. It would be favorable to investigate in the future which GAP enzyme can inhibit Rab2 specifically and to use it as a potential drug target.

## Funding

Supported by the ÚNKP-19-3 and ÚNKP-20-5 New National Excellence Program of the Ministry for Innovation and Technology.

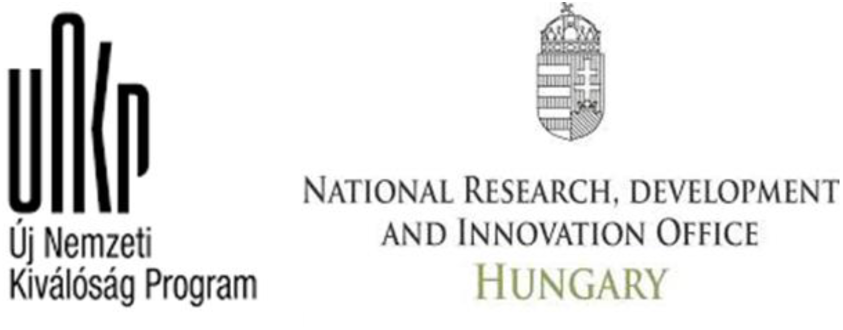

## Notes

### Competing Interest Statement

The authors have declared no competing interest.

